# Experimental determination of codon usage-dependent selective pressure on high copy-number genes in *Saccharomyces cerevisiae*

**DOI:** 10.1101/358259

**Authors:** Lyne Jossé, Tarun Singh, Tobias von der Haar

## Abstract

One of the central hypotheses in the theory of codon usage evolution is that in highly expressed genes particular codon usage patterns arise because they facilitate efficient gene expression and are thus selected for in evolution. Here we use plasmid copy number assays and growth rate measurements to explore details of the relationship between codon usage, gene expression level, and selective pressure in *Saccharomyces cerevisiae*. We find that when high expression levels are required optimal codon usage is beneficial and provides a fitness advantage, consistent with evolutionary theory. However, when high expression levels are not required, optimal codon usage is surprisingly and strongly selected against. We show that this selection acts at the level of protein synthesis, and we exclude a number of molecular mechanisms as the source for this negative selective pressure including nutrient and ribosome limitations and proteotoxicity effects. These findings inform our understanding of the evolution of codon usage bias, as well as the design of recombinant protein expression systems.

## Introduction

Because the genetic code uses 64 codons to encode 20 amino acids (Crick, Barnett, Brenner, & Watts-Tobin, 1961), most amino acids can be encoded by multiple synonymous codons. In all organisms where codon usage has been investigated, synonymous codons are not used randomly but some codons are preferred over others (see eg Behura & Severson, 2013) for a recent review). Correlations between biased codon usage and a number of other parameters have been detected including cellular tRNA content (Ikemura, 1982), translational efficiency (Sharp & Li, 1986), translational accuracy (Zhou, Weems, & Wilke, 2009), RNA structure (Hartl, Moriyama, & Sawyer, 1994), protein structure (Oresic, Dehn, Korenblum, & Shalloway, 2003), genomic GC content (Comeron, Kreitman, & Aguade, 1999), recombination (Comeron et al., 1999), splicing (Chamary & Hurst, 2005), and gene conversion rates (Galtier, 2003).

Translational efficiency has gained experimental support as one causative parameter that can lead to codon usage bias (Carlini & Stephan, 2003; Chu et al., 2014; Hense et al., 2010; Zhou et al., 2013). Since synonymous mutations can have substantial effects on expressed protein levels, selection for high gene expression levels will favour codon usage patterns compatible with such high levels, but will avoid patterns that restrict attainable expression levels. However, the exact relationship between codon usage, protein expression, other associated parameters and selective pressure are still unclear.

One issue is that selective pressure is difficult to measure directly as an experimental parameter. Moriya *et al.* used copy number variations in high copy number vectors in *Saccharomyces cerevisiae* to estimate the direction and extent of selective pressure when these vectors contained different genes (Makanae, Kintaka, Makino, Kitano, & Moriya, 2013; Moriya, Shimizu-Yoshida, & Kitano, 2006). While natural 2 micron circles employ a sophisticated system to stabilise copy numbers (McQuaid et al., 2017), yeast cloning vectors containing solely the 2 micron origin of replication exhibit strong copy number variations and appear to be inherited in a stochastic fashion (Moriya et al., 2006). In cases where such vectors contain genes that affect the growth rate of the cell in a copy-number dependent fashion, cells with reduced or increased copy number (depending on whether the gene in question has a negative or positive effect on growth rate) have a growth advantage. Since such cells are also more likely to produce daughter cells with copy numbers that are lower/ higher than the population average, the average copy number per cell changes until a new balance is achieved between the effect of the gene on growth rates, the metabolic cost of maintaining high numbers of vectors in the cell, and the ability to successfully inherit the vector and its selectable marker to the majority of daughter cells.

Here, we introduce reporter genes with differing codon usage into 2 micron cloning vectors and study the effect of varying codon usage on the high-copy number vector content and growth rate of transformed yeast cells. Surprisingly, our results suggest that codon optimisation is selected against unless high level expression of the gene provides a competitive growth advantage.

## Results

We previously described a dual luciferase reporter system allowing us to assess the effect of codon usage variation on Firefly luciferase expression levels (Chu et al., 2014; Chu, Barnes, & von der Haar, 2011). The original reporter system was implemented using centromeric (low copy number) vectors, which contained three different Firefly luciferase variants encoded only by unfavourable codons (minFLuc), by the wild-type gene from *Photinus pyralis* which has random codon usage from the point of view of yeast (staFLuc), or by the fully codon-adapted gene (maxFLuc). All vectors also contained an invariant *Renilla* luciferase gene for normalisation. For this vector system we had demonstrated that the expression ratio measured using dual luciferase assays corresponds well to the expression ratio measured by western blotting (Chu et al., 2011). We were therefore surprised to observe that, when the same expression system was implemented in multi-copy vectors based on the 2 micron origin of replication, the codon-dependent variation in expression levels was indistinguishable from the centromeric system when measured in dual luciferase assays, but differed markedly when assessed using western blots (figure 1). Note that all experiments shown in the current study were conducted with cytoplasmic firefly luciferase variants which contain deletions of the last three amino acids to remove a peroxisomal targeting signal (Chu et al., 2014).

**Figure 1.**
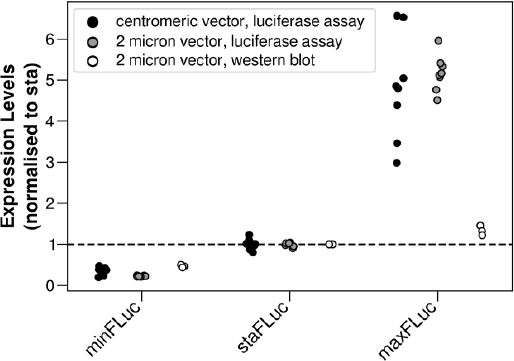
Expression levels of three codon usage variants of firefly luciferase. Expression levels were assessed either by dual luciferase assay and are shown as the ratio of expression between the variant firefly luciferases and an invariant *Renilla* luciferase gene also contained in the expression vectors (black and grey dots), or were assessed by western blotting in which case they are shown as absolute firefly expression levels (not normalised to *Renilla* levels).

We reasoned that our results could be explained if plasmid copy numbers changed substantially between expression plasmids containing the different codon usage variants. The dual luciferase assay measures expression normalised per plasmid, whereas the western blotting approach measures total numbers per cell, so that changes in plasmid copy number would only become visible in the latter assay. To test this assumption directly we measured the plasmid copy number for the different luciferase variants using a qPCR approach, and found that introduction of poorly adapted, wild type, and strongly codon adapted sequences sequentially reduced the steady-state plasmid copy number over a fourfold range compared to the vector control (a plasmid which only expresses *Renilla* luciferase but no firefly luciferase, figure 2b). This implies that expression of a strongly codon-adapted gene is less favourable and more strongly selected against than expression of weakly codon adapted genes. This effect is also visible at the levels of the growth rates of the respective transformants, which become successively reduced with increasing codon adaptation of the recombinant genes (figure 2c). This reduced growth rate appears due to lower cell division rates, as we do not detect any changes in the proportion of viable cells (data not shown).

**Figure 2.**
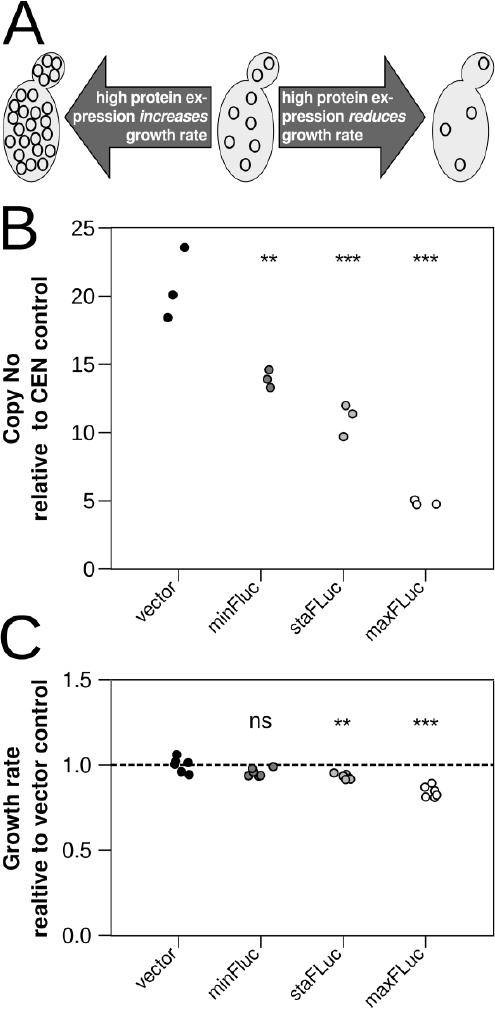
The expression of different firefly luciferase variants generates distinct fitness defects. **A,** the steady state copy number of 2 micron plasmids is modulated by the genes they contain. **B,** increasingly optimised codon usage leads to a decrease in steady state plasmid copy numbers as assessed by real time PCR. **C,** increasingly optimised codon usage in the recombinant luciferase gene leads to a decrease in growth rate. In both panels B and C, “vector” refers to a vector expressing only *Renilla* luciferase without a firefly luciferase gene. Statistical significance of results was assessed by ANOVA and Tukeys’ HSD test. Significance to the vector control is indicated as: ns, p>0.05; **, p<0.01; ***, p<0.001.

To test whether selection against codon optimised genes in this system arose at the level of protein synthesis, we introduced a point mutation into the codon adapted maxFLuc construct that resulted in a premature stop signal at the fifth codon of the open reading frame (maxFLuc K5X). This prevents almost all of the translational activity on the mRNA, without significantly affecting the overall sequence or nucleotide composition of the construct nor the transcriptional activity of the promoter. The K5X mutation substantially rescued both the reduction in plasmid copy number and the reduction in growth rate observed with the translated maxFLuc construct (figure 3). Thus, the major effect on fitness requires ongoing translation of the full-length ORF.

**Figure 3.**
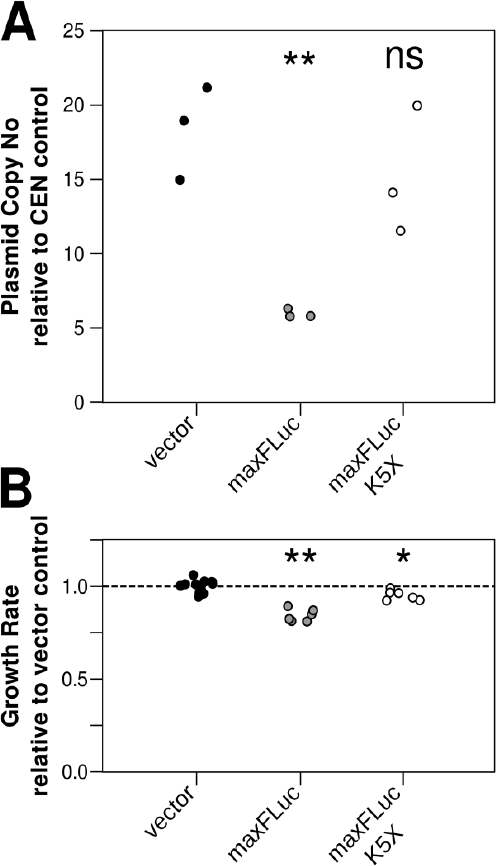
Fitness defects associated with the expression of codon optimised firefly luciferase are dependent on translation of the gene. **A,** plasmid copy numbers of a vector containing the codon optimised gene, the same gene with a nonsense mutation in the fifth codon, or the vector control are indicated. **B,** growth rate defects of the same strains as in panel A. Statistical significance of results was assessed by ANOVA and Tukeys’ HSD test. Significance to the vector control is indicated as: ns, p>0.05; *, p<0.05; **, p<0.01.

We explored a number of potential mechanisms which by our reasoning could have caused the selection against codon optimisation. We determined that the three constructs produced approximately 2.1×10^−14^, 4.5×10^−14^ and 5.9×10^−14^ g firefly luciferase protein per cell (figure 4A). In other words, the highest expressing construct produced just over 1% of the 5×10^−12^ g of protein contained in a haploid yeast cell (von der Haar, 2007). Most of the resources required for protein synthesis, including amino acids and energy, should scale linearly with the amount of protein made, and a 1-2% increase in resource usage would be unlikely to cause a 10-20% reduction in growth rate. Consistently, we do not observe any change in the effect of maxFLuc expression on the growth rate if we double the concentration of carbon, nitrogen, or amino acids in the medium (figure 4B).

**Figure 4.**
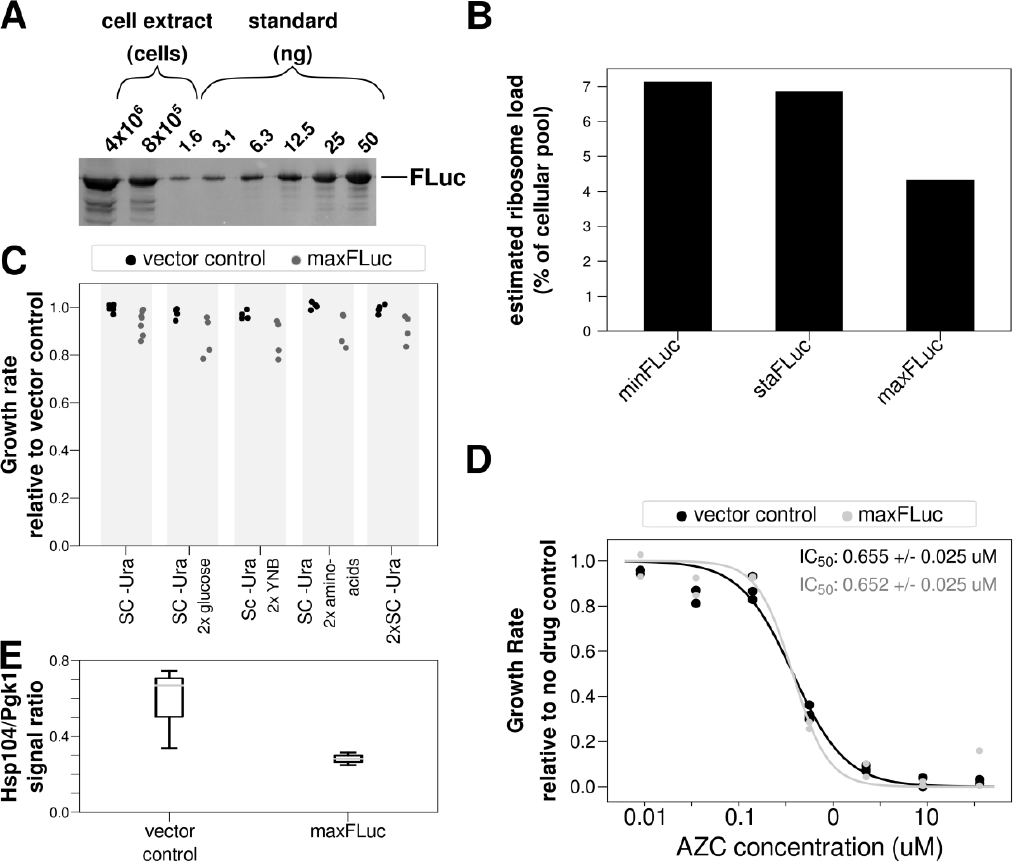
Exploring the mechanistic origin of fitness defects associated with the expression of a codon-optimised gene. **A,** the number of protein molecules per cell expressed from the staFLuc gene was determined by comparing western blot signals of recombinant luciferase with signals derived from controlled numbers of cells. **B,** estimated numbers of ribosomes required at steady state for translation of the different codon variants (see text for explanation). **C,** growth rate defects associated with expression of the codon optimised firefly luciferase gene are not altered in media supplemented with different nutrients. **D,** sensitivity of yeast strains containing the codon optimised firefly luciferase gene or a vector control to the protein unfolding drug azetidine-2-carboxylic acid (AZC). **E,** expression levels of the molecular chaperone Hsp104 (relative to the loading control Pgk1).

We and others previously predicted that, under fast growth conditions, providing ribosomes for the expression of additional proteins could only be accommodated at the cost of reduced growth (Chu et al., 2011; Chu & von der Haar, 2012; Shah, Ding, Niemczyk, Kudla, & Plotkin, 2013). We therefore estimated the number of ribosomes required to produce the observed protein levels from the three FLuc variants. This number essentially scales with the number of proteins produced per unit time, and inversely scales with the time spent to complete translation of the ORF (ie more ribosomes are needed to make more protein at the same speed, but less ribosomes are needed to make the same amount of protein at higher speed). In the case of our FLuc constructs, the increased speed of translation outweighs the more frequent initiation events for the maxFLuc construct, resulting in a predicted net decrease of ribosome usage with increasing codon optimisation (figure 4C). This decrease in ribosome usage for codon optimised genes is expected, and is thought to be one of the genome-wide drivers for codon optimisation as this maximises cell-wide ribosome availability (Shah & Gilchrist, 2011). In the context of our experiments, it rules out that ribosome shortages explain the selective pressure against codon optimised constructs. Lastly, we reasoned that simple proteotoxic effects might explain our results. However, the responsiveness of a maxFLuc expressing construct to the proteotoxic drug AZC (Trotter, Berenfeld, Krause, Petsko, & Gray, 2001) is unaltered (figure 4D), even though we would expect that any pre-existing proteotoxicity should increase the sensitivity to this drug. We also did not detect any significant increases in the expression of the disaggregase Hsp104, which is usually induced under conditions of proteotoxic stress (figure 4E, Bösl, Grimminger, & Walter, 2006).

In summary, although we clearly observe negative selective pressure against codon-optimised firefly luciferase, our experiments do not offer a simple explanation for this effect.

While we have no immediate mechanistic explanation for the strong negative selective pressure arising from codon optimisation in our construct, selection against the optimisation of a gene that is of no benefit to the cell makes some sense from an evolutionary perspective. To test how the dynamics of negative and positive selection interplay in the case of proteins that are of benefit to the cell, we repeated the FLuc expression experiments with the *HIS3* gene, a natural yeast gene which catalyses the 6^th^ step of the histidine biosynthesis pathway (Struhl & Davis, 1977). The yeast strain used in our experiments carries a chromosomal *his3* deletion and can only grow in the absence of histidine if the gene is provided extrachromosomally.

We determined the growth rates of yeast strains containing poorly adapted, wild-type, and well adapted *HIS3* genes under three different growth conditions. In the presence of histidine in the growth medium, the gene is not beneficial to the cell, similar to the FLuc genes in the experiments described above. In contrast, in the absence of histidine *HIS3* is essential, although it is not required at high expression levels (Chu et al., 2014). Lastly, in the absence of histidine and the presence of a competitive inhibitor of His3 enzymatic activity, 3-aminotriazole (3-AT), high levels of His3 protein are required to overcome the inhibitor (Durfee et al., 1993).

When His3 is not required, we observe a similar pattern of consecutively reduced fitness as for the FLuc genes (figure 5, upper panel). This confirms that the fitness effects observed upon high level FLuc expression are not specific to heterologous proteins and hold also upon high-level expression of a homologous gene. When His3 activity is required, the effect of the different constructs on growth rates is quantitatively changed and initially becomes statistically insignificant (figure 5, middle panel). When very high His3 levels are required in the presence of 3-AT, the codon-optimised gene confers a clear fitness advantage over the less optimised genes (figure 5, bottom panel). Links between preferred codon usage and the requirement for high expression levels have previously been established largely by way of correlation, our observations provide experimental confirmation of this notion.

**Figure 5.**
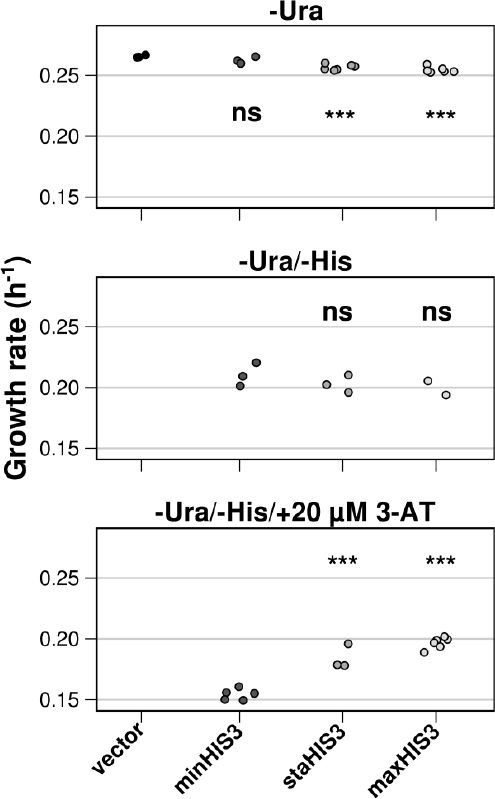
The interplay between required gene expression levels and fitness effects arising from codon optimisation. Growth rates of strains with a chromosomal deletion of the *HIS3* gene but containing codon variants of the same gene on 2 micron plasmids are shown. The plasmid borne-gene is expected to be superfluous in -Ura medium as this contains histidine, essential but required at low expression levels in -Ura/-His medium; and essential and required at high expression levels in the same medium containing 3-aminotriazole (3-AT), a competitive inhibitor of His3 enzymatic activity. Statistical significance of results was assessed by ANOVA and Tukeys’ HSD test. Significance to the vector control (top panel) or to the minHIS3 strains (middle and bottom panels) is indicated as: ns, p>0.05; ***, p<0.001.

To ask whether the effect of codon usage on selective pressure is quantitatively determined by basic yeast physiology, or is dependent on the genetic background of the strain in question, we compared the growth rates of a number of genetically tractable strains from the *Saccharomyces* genome resequencing project (SGRP, Cubillos, Louis, & Liti, 2009) in the presence and absence of the maxFLuc gene (figure 6). We found that the proportional reduction in growth rate upon expression of the gene ranged from 10% in the mildest affected strains, to 50% in the worst affected strains. Although many of the SGRP strains flocculate strongly and determination of growth rates for these strains is less accurate than for BY4741, this wide range indicates that the genetic make-up of strains strongly modulates the correspondence between codon usage and selective pressure.

**Figure 6.**
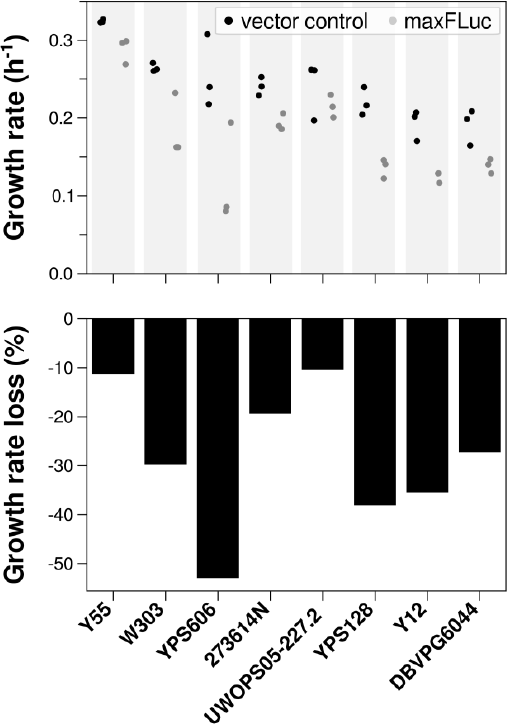
Fitness defects caused by expression of codon optimised firefly luciferase are strain dependent. Top, growth rates of strains containing the maxFLuc vector (grey dots) or a control vector (black dots). Strains are shown in descending order of average growth rate when containing the vector control construct. Bottom, average growth rate loss in % for strains containing the firefly luciferase gene compared to the vector control.

## Discussion

In interpreting our results, it is useful to define in how far plasmid copy number changes reflect selection mechanisms applying during the actual evolution of organisms. Codon usage is thought to have effects at both the individual gene level and at genome-wide levels. Effects at individual gene level include permitting high translational activity or conversely restricting translational activity to low levels (Tarrant & von der Haar (2014) and references therein), as well as effects on RNA stability (Presnyak et al., 2015) and transcriptional activity (Zhou et al., 2016). Genome-wide effects include compliance with particular GC bias (Palidwor, Perkins, & Xia, 2010), optimisation of ribosome usage by reducing the average dwell time of ribosomes on mRNAs (Chu & von der Haar, 2012; Shah et al., 2013) and other effects (Plotkin & Kudla, 2011). Genes contained in high copy number plasmids are in principle subject to effects at both levels, but because changing the sequence of such a plasmid is equivalent to changing the sequence of multiple genes simultaneously in the genome, we expect the selective forces acting on this system to be more similar to genome-wide selective forces than would be the case in a single copy plasmid.

The observation of particularly strong codon usage bias in highly expressed genes (Sharp, Tuohy, & Mosurski, 1986) implies that highly expressed genes show such bias, but also that low-expressed genes do not. It is thought that the absence of bias in genes with low expression levels arises because codon usage is at equilibrium, rather than being selected against (Sharp, Emery, & Zeng, 2010). Interestingly, our data suggest that codon bias can be selected against in genes with low expression levels. Moreover, the successive reduction in plasmid copy numbers with increasing codon optimisation (figure 2) suggests that negative selection is not restricted to extreme codon usage but applies to genes with close to equilibrium codon usage. Consistent with this notion, statistical enrichment of non-preferred codons in low expressed genes has been shown in some studies (Neafsey & Galagan, 2007).

The principal underlying assumption for explaining codon usage bias in highly expressed genes, that codon usage is selected for when high expression levels are required, has widespread support and is generally accepted. However, to our knowledge this has never been directly experimentally tested. By linking codon usage variants of the endogenous yeast *HIS3* gene to growth conditions where this gene is required at different expression levels, we provide an experimental test for this hypothesis. Our findings are entirely consistent with the prevailing hypothesis.

While our results are informative for understanding basic evolutionary mechanisms, they also provide useful information for the design of recombinant protein expression systems. 2 micron vector systems are used for the production of various yeast-derived biopharmaceuticals (Finnis et al., 2010; Gerngross, 2004; Thim et al., 1986). If plasmid copy numbers in these cases are subject to the same effects we describe here, productivity could be boosted by introducing measures that stabilise plasmid copy number, although this could also produce adverse effects by increasing the selection for non-expressing mutants. The observation of substantial strain variability in the effect of recombinant gene expression on growth rates (figure 6) also indicates that the genetic variability of yeast could be harnessed for balancing these effects in order to stabilise high level expression systems.

## Materials and Methods

*Strains, plasmids and media.* The main yeast strain used was BY4741 (Mat**a** *ura3Δ0 his3Δ0 leu2Δ0 met15Δ0*, Brachmann et al., 1998). Genetically tractable strains from the SGRP collection (Cubillos et al., 2009) were obtained from the National Collection of Yeast Cultures (NCYC, UK, haploid Mat**a** strains from SGRP strain set 2). Luciferase expression vectors were generated by transferring *XmaI/EcoRI* RLuc and *BamHI/SalI* CFLuc fragments from the centromeric vectors described in (Chu et al., 2014) into pBEVY-U (Miller, Martinat, & Hyman, 1998). *HIS3* expression vectors were generated by cloning the C-terminally HA-tagged *HIS3* variants described in (Chu et al., 2014) as *BamHI/PstI* fragments into the same vector. Plasmid sequences, maps, and the plasmids themselves are available through Addgene (table 1). Yeast strains were transformed as described (Gietz & Schiestl, 2007) and grown in SC -Ura medium (2% glucose, 0.67% Yeast Nitrogen Base without amino acids [BD, UK], and Kaiser Synthetic Complete Drop-Out Mixture lacking uracil [Formedium, UK] as indicated in the manufacturer’s instructions).

**Table 1.**
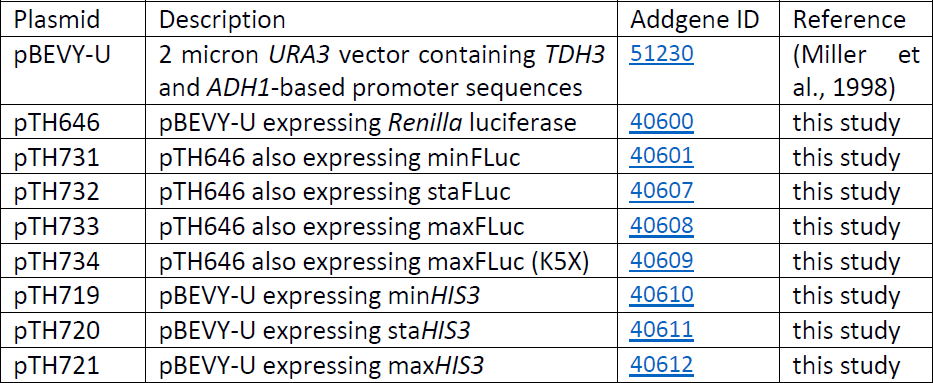
Plasmids used in this study.

*Growth rate measurements.* Growth rates were measured in 24- or 48-well cell culture plates using automated plate readers. 1 ml medium per well (0.5 ml for 48-well plates) were inoculated with material from transformed yeast colonies grown on selective agar plates and grown overnight in a standard shaking incubator. Following overnight growth, 1 ml or 0.5 ml of fresh medium per well contained in a new plate was inoculated with 10 µl or 5 µl of culture from the overnight plate. Plates were then incubated in Spectrostar Nano plate readers (BMG Lab Tech, UK) and incubated at 30°C under constant shaking, with automated oD measurements every 30 minutes until the culture had reached stationary phase. To analyse the resulting data, logarithmic curves were fitted to seven-time-point windows along the entire incubation time, and the highest growth rate returned from this fitting exercise was reported as the maximum logarithmic growth rate for each well.

*qPCR assays.* These were adapted from the procedure described in (Moriya et al., 2006). Briefly, 2 oD_600_ units of cells were resuspended in 850 µl of 1.2 M sorbitol, 100 µl of 200 mM sodium phosphate buffer (pH 7.2), and 50 µl lyticase (Sigma Aldrich, UK, L2524, resuspended in sodium phosphate buffer at 5 units/ µl). The suspension was incubated at 37°C for 30 minutes, and then subjected to 3 cycles of incubation for ten minutes at 95°C followed by incubation for fifteen minutes at -80°. The extract was clarified by centrifugation in a microcentrifuge at 13,000 rpm for 5 minutes, and 0.1 µl of the supernatant were used as input for the qPCR reactions.

For the latter, we used primers against the *URA3* marker gene on the pBEVY-U vectors (qURA3f, AGCAGAATTGTCATGCAAGG, qURA3r, TTCCACCCATGTCTCTTTGA) and against the *LEU3* gene for standardisation (Chu et al., 2014). qPCR reactions were assembled using QuantiFast SYBR Green PCR Kits (QIAgen, UK) according to the manufacturer’s instructions, and reactions were run using a two-step amplification protocol. Ct values were determined in the logarithmic amplification range and converted to fold change values as described (Pfaffl, 2001). All reported data were determined using triplicate biological replicates, each assayed in technical duplicate.

*Protein extraction and western blotting* were conducted as described (von der Haar, 2007), using antibodies from Sigma-Aldrich, UK (anti-firefly luciferase, L0159, anti-HA tag, H6908, and peroxidase-labelled anti-rabbit IgG, A9169). Anti-Hsp104 antibodies were as described (Adam, Jossé, & Tuite, 2017).

*Data analysis and statistics.* All data analysis scripts used to prepare individual figures are available for download from Github^1^ or for execution online from MyBinder^2^. Statistical significance between samples was tested using ANOVA followed by Tukey’s Honestly Significant Difference test.

## Acknowledgements

This work was funded by the Biotechnology and Biological Sciences Research Council (UK, BBSRC) through grant BB/I010351/1 and by the Royal Society (UK) through grant RG090115.

## Footnotes

1 https://github.com/tobiasvonderhaar/codonselection

2 https://mybinder.org/v2/gh/tobiasvonderhaar/codonselection/master

## References

Adam, I., Jossé, L., & Tuite, M. F. (2017). Human TorsinA can function in the yeast cytosol as a molecular chaperone. Biochemical Journal, 474(20), 3439–3454. https://doi.org/10.1042/BCJ20170395

Behura, S. K., & Severson, D. W. (2013). Codon usage bias: causative factors, quantification methods and genome-wide patterns: with emphasis on insect genomes. Biological Reviews of the Cambridge Philosophical Society, 88(1), 49–61. https://doi.org/10.1111/j.1469-185X.2012.00242.x

Bösl, B., Grimminger, V., & Walter, S. (2006). The molecular chaperone Hsp104—A molecular machine for protein disaggregation. Journal of Structural Biology, 156(1), 139–148. https://doi.org/10.1016/j.jsb.2006.02.004

Brachmann, C. B., Davies, A., Cost, G. J., Caputo, E., Li, J., Hieter, P., & Boeke, J. D. (1998). Designer deletion strains derived from Saccharomyces cerevisiae S288C: a useful set of strains and plasmids for PCR-mediated gene disruption and other applications. Yeast, 14(2), 115–132. https://doi.org/10.1002/(SICI)1097-0061(19980130)14:2<115::AIDYEA204>3.0.CO;2-2

Carlini, D. B., & Stephan, W. (2003). In Vivo Introduction of Unpreferred Synonymous Codons Into the Drosophila Adh Gene Results in Reduced Levels of ADH Protein. Genetics, 163, 239–243.

Chamary, J.-V., & Hurst, L. D. (2005). Biased codon usage near intron-exon junctions: selection on splicing enhancers, splice-site recognition or something else? Trends in Genetics: TIG, 21(5), 256–9. https://doi.org/10.1016/j.tig.2005.03.001

Chu, D., Barnes, D. J., & von der Haar, T. (2011). The role of tRNA and ribosome competition in coupling the expression of different mRNAs in Saccharomyces cerevisiae. Nucleic Acids Research, 39(15), 6705–6714. https://doi.org/10.1093/nar/gkr300

Chu, D., Kazana, E., Bellanger, N., Singh, T., Tuite, M. F., & von der Haar, T. (2014). Translation elongation can control translation initiation on eukaryotic mRNAs. The EMBO Journal, 33(1), 21–34. https://doi.org/10.1002/embj.201385651

Chu, D., & von der Haar, T. (2012). The architecture of eukaryotic translation. Nucleic Acids Research, 40(20), 10098–10106. https://doi.org/10.1093/nar/gks825

Comeron, J. M., Kreitman, M., & Aguade, M. (1999). Natural Selection on Synonymous Sites Is Correlated With Gene Length and Recombination in Drosophila. Genetics, 151, 239–249.

Crick, F. H., Barnett, L., Brenner, S., & Watts-Tobin, R. J. (1961). General nature of the genetic code for proteins. Nature, 192, 1227–1732. Retrieved from http://www.ncbi.nlm.nih.gov/pubmed/13882203

Cubillos, F. a, Louis,, E. J., & Liti,, G. (2009). Generation of a large set of genetically tractable haploid and diploid Saccharomyces strains. FEMS Yeast Research, 9(8), 1217–1225. https://doi.org/10.1111/j.1567-1364.2009.00583.x

Durfee, T., Becherer, K., Chen, P. L., Yeh, S. H., Yang, Y., Kilburn, A. E., … Elledge, S. J. (1993). The retinoblastoma protein associates with the protein phosphatase type 1 catalytic subunit. Genes & Development, 7(4), 555–569. Retrieved from http://www.ncbi.nlm.nih.gov/pubmed/8384581

Finnis, C. J. A., Payne, T., Hay, J., Dodsworth, N., Wilkinson, D., Morton, P., … Sleep, D. (2010). High-level production of animal-free recombinant transferrin from Saccharomyces cerevisiae. Microbial Cell Factories, 9, 87. https://doi.org/10.1186/1475-2859-9-87

Galtier, N. (2003). Gene conversion drives GC content evolution in mammalian histones. Trends in Genetics, 19(2), 65–68. Retrieved from http://www.ncbi.nlm.nih.gov/pubmed/12547511

Gerngross, T. U. (2004). Advances in the production of human therapeutic proteins in yeasts and filamentous fungi. Nature Biotechnology, 22(11), 1409–1414. https://doi.org/10.1038/nbt1028

Gietz, R. D., & Schiestl, R. H. (2007). Large-scale high-efficiency yeast transformation using the LiAc/SS carrier DNA/PEG method. Nature Protocols, 2(1), 38–41. https://doi.org/10.1038/nprot.2007.15

Hartl, D. L., Moriyama, E. N., & Sawyer, S. A. (1994). Selection intensity for codon bias. Genetics, 138(1), 227–34. Retrieved from http://www.ncbi.nlm.nih.gov/pubmed/8001789

Hense, W., Anderson, N., Hutter, S., Stephan, W., Parsch, J., & Carlini, D. B. (2010). Experimentally increased codon bias in the Drosophila Adh gene leads to an increase in larval, but not adult, alcohol dehydrogenase activity. Genetics, 184(2), 547–55. https://doi.org/10.1534/genetics.109.111294

Ikemura, T. (1982). Correlation between the abundance of yeast transfer RNAs and the occurrence of the respective codons in protein genes. Differences in synonymous codon choice patterns of yeast and Escherichia coli with reference to the abundance of isoaccepting transfer R. Journal of Molecular Biology, 158(4), 573–597. https://doi.org/10.1016/0022-2836(82)90250-9

Makanae, K., Kintaka, R., Makino, T., Kitano, H., & Moriya, H. (2013). Identification of dosage-sensitive genes in Saccharomyces cerevisiae using the genetic tug-of-war method. Genome Research, 23(2), 300–311. https://doi.org/10.1101/gr.146662.112

McQuaid, M. E., Pinder, J. B., Arumuggam, N., Lacoste, J. S. C., Chew, J. S. K., & Dobson, M. J. (2017). The yeast 2-μm plasmid Raf protein contributes to plasmid inheritance by stabilizing the Rep1 and Rep2 partitioning proteins. Nucleic Acids Research, 45(18), 10518–10533. https://doi.org/10.1093/nar/gkx703

Miller, C. A. I., Martinat, M. A., & Hyman, L. E. (1998). Assessment of aryl hydrocarbon receptor complex interactions using pBEVY plasmids: expression vectors with bidirectional promoters for use in Saccharomyces cerevisiae. Nucleic Acids Research, 26(15), 3577–3583.

Moriya, H., Shimizu-Yoshida, Y., & Kitano, H. (2006). In vivo robustness analysis of cell division cycle genes in Saccharomyces cerevisiae. PLoS Genetics, 2(7), e111. https://doi.org/10.1371/journal.pgen.0020111

Neafsey, D. E., & Galagan, J. E. (2007). Positive selection for unpreferred codon usage in eukaryotic genomes. BMC Evolutionary Biology, 7(1), 119. Retrieved from http://www.pubmedcentral.nih.gov/articlerender.fcgi?artid=1936986&tool=pmcentrez&rendertype=abstract

Oresic, M., Dehn, M., Korenblum, D., & Shalloway, D. (2003). Tracing specific synonymous codon-secondary structure correlations through evolution. Journal of Molecular Evolution, 56(4), 473–484. https://doi.org/10.1007/s00239-002-2418-x

Palidwor, G. A., Perkins, T. J., & Xia, X. (2010). A General Model of Codon Bias Due to GC Mutational Bias. PLoS ONE, 5(10), e13431. https://doi.org/10.1371/journal.pone.0013431

Pfaffl, M. W. (2001). A new mathematical model for relative quantification in real-time RTPCR. Nucleic Acids Research, 29(9), e45. Retrieved from http://www.pubmedcentral.nih.gov/articlerender.fcgi?artid=55695&tool=pmcentrez&rendertype=abstract

Plotkin, J. B., & Kudla, G. (2011). Synonymous but not the same: the causes and consequences of codon bias. Nature Reviews Genetics, 12(1), 32–42. https://doi.org/10.1038/nrg2899

Presnyak, V., Alhusaini, N., Chen, Y.-H., Martin, S., Morris, N., Kline, N., … Coller, J. (2015). Codon Optimality Is a Major Determinant of mRNA Stability. Cell, 160(6), 1111–1124. https://doi.org/10.1016/j.cell.2015.02.029

Shah, P., Ding, Y., Niemczyk, M., Kudla, G., & Plotkin, J. B. (2013). Rate-Limiting Steps in Yeast Protein Translation. Cell, 153(7), 1589–1601. https://doi.org/10.1016/j.cell.2013.05.049

Shah, P., & Gilchrist, M. A. (2011). Explaining complex codon usage patterns with selection for translational efficiency, mutation bias, and genetic drift. Proceedings of the National Academy of Sciences of the United States of America, 108(25), 10231–10236. https://doi.org/10.1073/pnas.1016719108

Sharp, P. M., Emery, L. R., & Zeng, K. (2010). Forces that influence the evolution of codon bias. Philosophical Transactions of the Royal Society of London. Series B, Biological Sciences, 365(1544), 1203–1212. https://doi.org/10.1098/rstb.2009.0305

Sharp, P. M., & Li, W. H. (1986). An evolutionary perspective on synonymous codon usage in unicellular organisms. Journal of Molecular Evolution, 24(1–2), 28–38. Retrieved from http://www.ncbi.nlm.nih.gov/pubmed/3104616

Sharp, P. M., Tuohy, T. M. F., & Mosurski, K. R. (1986). Codon usage in yeast: cluster analysis clearly differentiates highly and lowly expressed genes. Nucleic Acids Research, 14(13), 5125–5143. https://doi.org/10.1093/nar/14.13.5125

Struhl, K., & Davis, R. W. (1977). Production of a functional eukaryotic enzyme in Escherichia coli: cloning and expression of the yeast structural gene for imidazole-glycerolphosphate dehydratase (his3). Proceedings of the National Academy of Sciences of the United States of America, 74(12), 5255–5259. Retrieved from http://www.ncbi.nlm.nih.gov/pubmed/341150

Tarrant, D., & Von Der Haar, T. (2014). Synonymous codons, ribosome speed, and eukaryotic gene expression regulation. Cellular and Molecular Life Sciences, 71(21), 4195–4206. https://doi.org/10.1007/s00018-014-1684-2

Thim, L., Hansen, M. T., Norris, K., Hoegh, I., Boel, E., Forstrom, J., … Fiil, N. P. (1986). Secretion and processing of insulin precursors in yeast. Proceedings of the National Academy of Sciences, 83(18), 6766–6770. https://doi.org/10.1073/pnas.83.18.6766

Trotter, E. W., Berenfeld, L., Krause, S. A., Petsko, G. A., & Gray, J. V. (2001). Protein misfolding and temperature up-shift cause G1 arrest via a common mechanism dependent on heat shock factor in Saccharomycescerevisiae. Proceedings of the National Academy of Sciences of the United States of America, 98(13), 7313–7318. https://doi.org/10.1073/pnas.121172998

von der Haar, T. (2007). Optimized protein extraction for quantitative proteomics of yeasts. PloS One, 2(10), e1078. https://doi.org/10.1371/journal.pone.0001078

Zhou, M., Guo, J., Cha, J., Chae, M., Chen, S., Barral, J. M., … Liu, Y. (2013). Non-optimal codon usage affects expression, structure and function of clock protein FRQ. Nature, 495(7439), 111–115. https://doi.org/10.1038/nature11833

Zhou, T., Weems, M., & Wilke, C. O. (2009). Translationally optimal codons associate with structurally sensitive sites in proteins. Molecular Biology and Evolution, 26(7), 1571–80. https://doi.org/10.1093/molbev/msp070

Zhou, Z., Dang, Y., Zhou, M., Li, L., Yu, C., Fu, J., … Liu, Y. (2016). Codon usage is an important determinant of gene expression levels largely through its effects on transcription. Proceedings of the National Academy of Sciences, 113(41), E6117–E6125. https://doi.org/10.1073/pnas.1606724113

